# Development of hybrid alphavirus-influenza A, B, and D pseudovirions for rapid quantification of neutralization antibodies and antiviral drugs

**DOI:** 10.64898/2026.03.03.709407

**Authors:** Brian Hetrick, Dongyang Yu, Eric Mazur, Kripa Giri, Feng Li, Dan Wang, Kylene Kehn-Hall, Yuntao Wu

## Abstract

The emergence and spread of highly pathogenic avian influenza H5N1 subtypes have raised global concerns due to their ability to cross species barriers and occasional spillover to humans. The viruses primarily infect wild birds and poultry, which have caused significant, sporadic outbreaks in mammals including dairy cattle. Influenza D virus is a recently identified influenza virus that mainly affects cattle with frequent spillover to other species such as swine. Despite the availability of poultry vaccines, there are no H5N1 and Influenza D vaccines for cattle or other potentially affected livestock. Given a history of frequent influenza pandemics originating from avian and mammalian hosts, there is an urgent need for enhanced surveillance, biosecurity, and the development of antivirals and vaccines. Here we describe the development of a novel hybrid alphavirus-influenza pseudovirion (Ha-IV), which is a non-replicating influenza virus-like particle composed of viral structural proteins and an RNA genome derived from a fast-expressing alphaviral vector. As a proof-of-concept, we assembled Ha-IV pseudoviruses based on influenza D and influenza A and B subtypes, and demonstrated their infectivity. In addition, we validated an influenza A pseudovirus based on the H5N1 clade 2.3.4.4b strain, A/Texas/37/2024, for rapid quantification of neutralization antibodies within 4 to 18 hours. Furthermore, we used the pseudovirus to quantify infected cow sera and performed a correlation study with the classic hemagglutinin inhibition assay (HIA). We demonstrate that the Ha-IAV pseudovirus-based assay is consistent with HIA in identifying protective antibody responses. Our results demonstrate that this new Ha-IV pseudovirus provides a rapid tool for quantifying the infectivity of emerging HA mutants and for assessing neutralizing antibody responses.

## 1. Introduction

Since 2021, highly pathogenic avian influenza (HPAI) H5N1 subtypes have been increasingly detected in poultry and wild birds [1]. Between 2020 and 2023, over 48 mammalian species across 26 countries were reported to be infected by H5N1 [2]. The U.S. reported its first human H5N1 clade 2.3.4.4b case in April 2024 when a dairy worker in Texas developed mild symptoms. This human infection coincided with unexpected cases of H5N1 clade 2.3.4.4b in dairy cows across multiple U.S. states, signaling a concerning trend of the virus crossing species barriers [3, 4]. Historically, influenza pandemics, originating from avian or mammal hosts, have occurred four times since the early 20th century, with the Spanish flu being the most severe. The ongoing evolution of influenza viruses and the recent mammalian infections have raised concerns about potential pandemics [5].

Influenza D virus (IDV) is a recently identified member of the Orthomyxoviridae, and was first detected in swine in a Oklahoma pig farm and subsequently identified in cattle across the USA [6]. Serological evidence of IDV infection has since been reported in a broad range of domestic and wild species, including camels, horses, small ruminants, feral and wild pigs, hedgehogs, giraffes, kangaroos, wallabies, and llamas [7]. Despite this unusually broad host range, which somehow resembles HPAI H5N1, IDV remains the least characterized influenza virus, and its epidemiology and zoonotic potential remain largely unresolved.

The CDC and WHO emphasize global surveillance and the monitoring of emerging influenza strains, including H5N1. The CDC’s pandemic risk assessment involves evaluating the threat posed by the H5N1 strain using the Influenza Risk Assessment Tool (IRAT). Given the H5N1 virus’s global spread among wild birds and mammals, there is an urgent need for enhanced surveillance and biosecurity to mitigate the risk of a potential pandemic. The hemagglutinin inhibition assay (HIA) is a well-established technique in influenza research, widely used as a gold standard assay for vaccine efficacy studies, seroepidemiological investigations, and the assessment of antigenic drift in influenza viruses. However, HIA has some limitations in quantifying viral infectivity and neutralizing antibodies; especially, the H5 viruses inherently have a varied ability to cause agglutination. For example, the assay does not accurately quantify the ability of different H5N1 variants to escape antibody neutralization or assess the efficiency of viral entry into host cells [8]. The standard for quantifying viral infectivity and neutralizing antibodies typically involves the use of infectious viruses or pseudoviruses. The use of HPAI such as H5N1 requires biosafety level 3 (BSL-3) facilities and practices, which are not widely available and are impractical for routine clinical testing. To overcome these challenges, pseudoviruses are increasingly being utilized in influenza research. Pseudoviruses can directly quantify the hemagglutinin (HA)-mediated viral entry and the neutralizing antibody titer of serum samples in a BSL-2 setting. The most common pseudotyped viruses for H5N1 research are based on the vesicular stomatitis virus (VSV) and the lentivirus. The Influenza H5N1 pseudotyped VSV platform consists of a replication-deficient reporter virus that lacks the major VSV glycoprotein. This system can produce a large quantity of pseudoviral particles at scale; however, it suffers from complex assembly schemes and is not as practical as pseudotyped lentiviruses. Pseudotyped lentiviruses are produced by co-transfection of plasmids expressing the lentivirus genome, the genes for lentiviral gag-pol structural proteins, and the HA gene from H5N1. Both VSV and lentivirus systems typically require a minimum of 24 to 48 hours to generate sufficient reporter signals for quantifying infection, making them challenging for large-scale clinical testing. Additionally, these pseudovirus systems only include the HA protein of H5N1, excluding other structural proteins such as neuraminidase (NA), matrix 1 (M1) and matrix 2 (M2). Thus, these pseudoviruses cannot be used to explore antivirals that impact influenza viral budding and other functional steps that are mediated by the viral structural proteins.

In this article, we describe the development of a novel influenza pseudovirus: the hybrid alphavirus-influenza pseudovirion (Ha-IV). Ha-IV consists of a viral particle formed from the structural proteins of influenza viruses: the HA, NA, M1 and M2, the nucleoprotein (NP), or the influenza D hemagglutinin-esterase-fusion (HEF). In addition, the particle encapsulates an RNA genome derived from a fast-expressing alphaviral vector for rapid reporter gene expression. For proof-of-concept, we assembled a pseudovirus based on the H5N1 clade 2.3.4.4b strain, A/Texas/37/2024, and validated this pseudovirus for the rapid quantification of neutralization antibodies and antiviral drugs.

## 2. Materials and Methods

### 2.1 Virus and viral particle assembly

The Influenza A structural protein expression vectors (pCMV-H5N1-HA, pCMV-H5N1-NA, pCMV-H5N1-NP, pCMV-H5N1-M1, pCMV-H5N1-M2) and the alphaviral vectors, alpha-PS5769-dsRED-Express2 or alpha-PS5769-Luc were provided by Virongy Biosciences Inc. Ha-IV particles were assembled by co-transfection of HEK293T cells with Influenza A virus (IAV) structural protein expression vectors (HA, NA, NP, M1 and M2) and the alphaviral dsRED-Express2 vector or alphaviral Luc vector as previously described [9]. For Ha-IDV assembly, pIDV-HEF was used to replace the pCMV-H5N1-HA and pCMV-H5N1-NA vectors. Particles were harvested at 48 hours post cotransfection, filtered through a 0.45 µm filter.

### 2.2 Viral infectivity assay

Ha-IV particles were used to infect HEK293T cells or A459 (ATCC) or A459(DC-SIGN) cells. Briefly, cells were seeded in 12-well plates (2 x 10^5^ cells per well), and infected for 4 - 24 hours at 37°C, and then observed for dsRED expression, or lysed in luciferase lysis and assay buffer (Virongy) for luciferase activity using GloMax Discover Microplate Reader (Promega).

### 2.3 Ha-IV neutralizing antibody assay

Neutralization assay was performed in 96-well plate. Ha-IAV(A/Texas/37/2024-H5N1)(Luc) particles (45 μl) were pre-incubated with 15 μl serially diluted goat anti-HA polyclonal serum (NR-10274) (BEI Resources, NIAID, NIH) or cow sera (provided by the Maxwell H. Gluck Equine Research Center, University of Kentucky, two uninfected, one from 7 days post-infection, and one from 10 days post-infection) for 10 minutes at 37°C, and then used to infect 15 μl HEK293T cells (5×10^4^ cells per well) for 18 hours at 37 °C. Cells were lysed in 7.5 μl Cell Lysis Buffer (Virongy) and 25 μl Luciferase Assay Buffer (Virongy) were added for luciferase assays using GloMax Discover Microplate Reader (Promega). The IC_50_ and neutralization curve was calculated using GraphPad PRISM software.

### 2.3. Hemagglutination and hemagglutination inhibition assays

Influenza A hemagglutination assay (HA) was performed in V-bottom shaped 96-well plate using 100 μl of turkey red blood cells (Lampire Biological Laboratories) suspended in PBS and mixed with 50 μl dilutions of the Ha-IVA (H5N1-A/Texas/37/2024-H5N1) pseudovirus particle. Plates were incubated at room temperature for approximately 30 minutes until the control red blood cells completely settle. HA titer was determined by the denominator of the last dilution which complete hemagglutination was seen. For hemagglutination inhibition assay (HIA), Ha-IAV(A/Texas/37/2024-H5N1)(Luc) particles (45 μl) were pre-incubated with 15 μl serially diluted cow sera (provided by the Maxwell H. Gluck Equine Research Center, University of Kentucky, two uninfected, one from 7 days post-infection, and one from 10 days post-infection) for 10 minutes at 37°C, and then mixed with 100 μl of turkey red blood cells in V-bottom shaped 96-well plate. Plates were incubated at room temperature for approximately 30 minutes until the control red blood cells completely settle. HIA titer was determined by the denominator of the last dilution which complete hemagglutination was seen.

### 2.4. Arbidol inhibition assay

Arbidol-inhibition assay was performed in 96-well plate. HEK293T cells (5×10^4^ cells per well in 15 μl) was pre-treated with 5 μl serially diluted Arbido (Tocris, dissolved in DMSO) for 1 hour, and then infected with Ha-IAV(A/Texas/37/2024-H5N1)(Luc) particles (45 μl) in the presence of Arbidol for 18 hours at 37 °C. Cells were lysed in 7.5 μl Cell Lysis Buffer (Virongy), and 25 μl Luciferase Assay Buffer (Virongy) were added for luciferase assays using GloMax Discover Microplate Reader (Promega). The IC_50_ and neutralization curve was calculated using GraphPad PRISM software.

### 2.5. Quantification and statistical analysis

Luciferase expression was quantified as the mean value of three luciferase assay readings, and the standard deviation (SD) values were calculated using Microsoft Excel. The neutralization activities of the antibodies were plotted and the IC_50_ values were calculated using GraphPad Prism 7.

## 3. Results

### 3.1. Design of the Ha-IV pseudoviruses

Previously, we developed a hybrid alphavirus-SARS-Cov-2 (Ha-CoV-2) pseudovirus for rapid quantification of neutralizing antibodies [9]. The pseudovirus consists of a virus-like particle assembled from the SARS-CoV-2 structural proteins, which encapsulate a modified self-amplifying RNA replicon derived from an attenuated, BSL-2-safe Semliki Forest virus (SFV). This RNA replicon consists of the 5’ untranslated region and open-reading frames coding for the SFV nonstructural proteins (nsP) 1-4 [10, 11], which allow for rapid amplification of the RNA in target cells. Previous studies have demonstrated that alphavirus replicon can quickly express genes to produce up to 200,000 copies of its RNA and large quantities of reporter proteins from its subgenomic promoter within hours after entering cells [11, 12]. This RNA replicon also contains a viral subgenomic RNA promoter that is used for the rapid expression of reporter genes (such as GFP or luciferase).

We used similar design of Ha-CoV-2 to develop the hybrid alphavirus-influenza (Ha-IV) pseudoviruses. As shown in **Figure 1**, the hybrid alphavirus-influenza A virus (Ha-IAV) and -influenza B virus (Ha-IBV) particles are assembled by cotransfection of HEK293T cells with DNA vectors expressing the structural proteins of IAV (HA, NA, M1, M2, and NP) and the genomic alphaviral RNA replicon (**Figure 1A**). For the assembly of the hybrid alphavirus-influenza D (Ha-IVD) particles, we replaced the influenza HA and NA expression vectors with a vector expressing the influenza D HEF protein (**Figure 1B**). Particles are harvested at 48 hours post co-transfection and tested for the ability to infect target cells.

**Figure 1.**
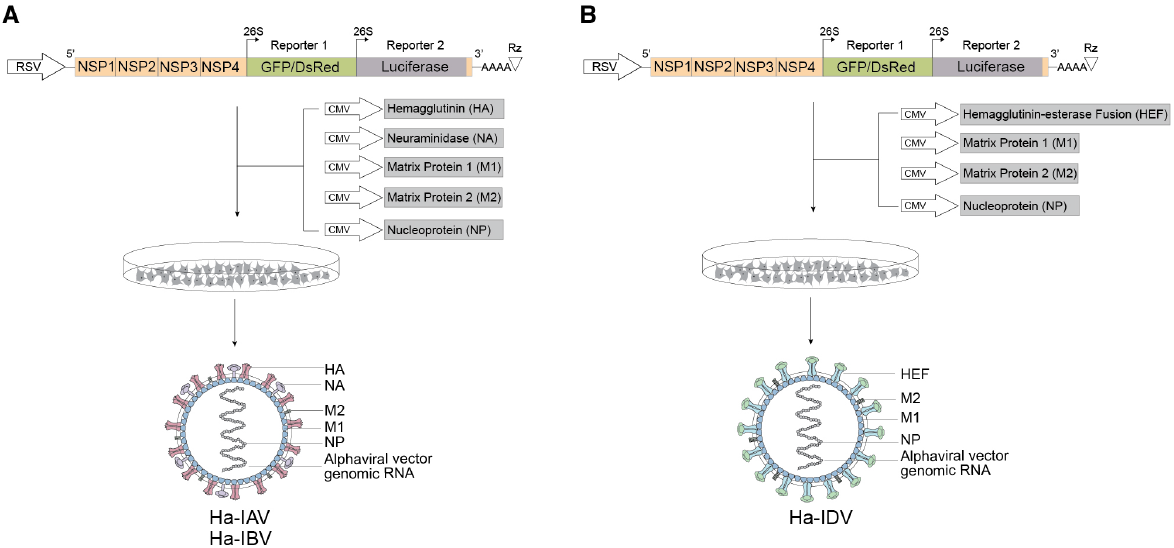
Illustration of the design and assembly of Ha-IV pseudoviral particles. (**A**) The design and assembly of Ha-IAV and Ha-IBV particles. The alphavirus-based genomic vector contains a RSV promoter that transcribes viral RNA genome that is packaged into Ha-IV particles. Shown are the 5’ untranslated region followed by the ORFs of the nonstructural proteins (nsp) 1-4 from SFV, viral subgenomic promoters for reporter expression, the 3’ untranslated region, and a poly(A) tail that is self-cleaved by the hepatitis delta virus ribozyme (RZ). To assemble viral particles, HEK293T cells were co-transfected with the SFV alphaviral vector and the vectors expressing the 5 structural proteins of influenza A (HA, NA, M1, M2, and NP). Viral particles in the supernatant were harvested at 48 hours. (**B)** The design and assembly of Ha-IDV particles. For particle assembly, the influenza D surface glycoprotein HEF was used to replace HA and NA.

### 3.2. Infectivity of Ha-IAV, Ha-IBV, and Ha-IDV pseudoviruses

To demonstrate the infectivity of the Ha-IAV pseudovirus, we assembled the particle based on the influenza A H5N1 clade 2.3.4.4b strain, A/Texas/37/2024. The structural protein genes are derived from the Genebank PP577943-PP577946 sequences. We also used the alphaviral vector RNA replicon to express the dsRED-Express2 reporter. Particles were assembled and then used to infect HEK293T target cells. As shown in **Figure 2A**, we observed dsRED-Express2 expression in target cells following viral infection. To further confirm the ability of the pseudovirus for rapid reporter expression, we constructed another reporter virus expressing the firefly luciferase reporter (Luc). We conducted a time-course experiment to monitor reporter expression in cells following infection. Intracellular luciferase activities were quantified at 4, 12, and 24 hours post-infection. As shown in **Figure 2B**, luciferase activity was detected as early as 4 hours post infection and increased with time, reaching the highest level at 24 hours in this time-course experiment.

**Figure 2.**
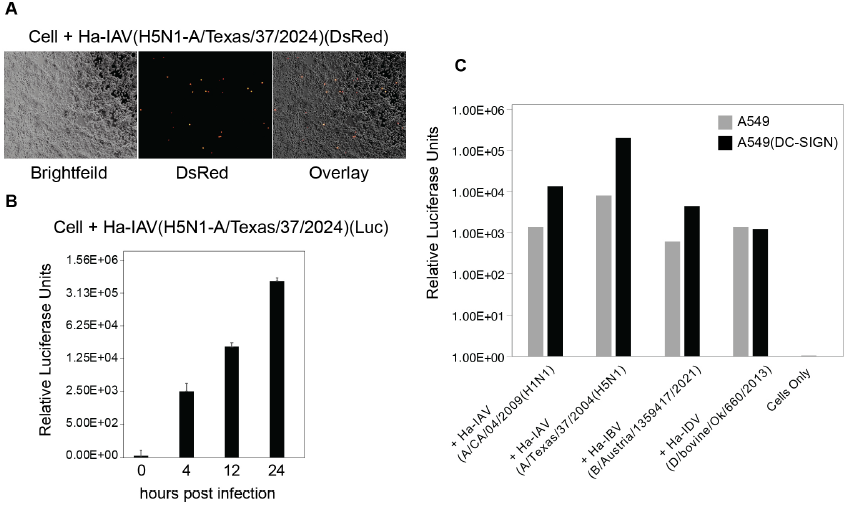
Validation of Ha-IV. (**A**) Pseudoviral particle, Ha-IAV(A/Texas/37/2024-H5N1)(DsRed) carrying the DsRed-Express2 reporter gene are assembled and used to infect HEK293T cells. DsRed-Express2 expression was observed 24 hours post infection. (**B**) Rapid luciferase (Luc) reporter expression following Ha-IAV(A/Texas/37/2024-H5N1)(Luc) infection. Cells were infected with the Luc reporter pseudovirus for 4 to 24 hours, and then lysed and analyzed for Luc expression. Time “0” is uninfected cells, hours post infection begin at the time of virus addition. Luciferase assays were performed in technical triplicates, and the mean and standard deviation (SD) are shown. (**C**) Infectivity of Ha-IAV, Ha-IBV, and Ha-IDV pseudoviruses. Pseudoviral particles carrying the Luc reporter were assembled and used to infect A459 or A549(DC-SIGN) target cells. Luc reporter expression following Ha-IAV(A/California/04/2009-H1N1)(Luc), Ha-IAV(A/Texas/37/2004-H5N1)(Luc), Ha-IBV(B/Austria/1359417/2021(Luc), or Ha-IDV(D/bovine/Oklahoma/660/2013)(Luc) infection was quantified. Cells were infected with the Luc reporter pseudoviruses for 18 hours, lysed, and analyzed for Luc expression.

We also assembled additional Ha-IAV pseudoviruses based on other subtypes such as the influenza A California/04/2009 (H1N1), which was first identified in Southern California during the 2009 H1N1 pandemic [13]. The virus is a swine-origin virus and is frequently used in research for vaccine development and pathogenicity studies [14]. In addition, we assembled a Ha-IBV and a Ha-IDV pseudovirus based on the influenza B Austria/1359417/2021 [15], and a recently discovered influenza D virus (influenza D/bovine/Oklahoma/660/2013) [6], respectively. The infectivity of these pseudoviruses were confirmed by infecting the human alveolar basal epithelial adenocarcinoma cell line A549 (**Figure 2C**). Following infection for 18 hours, a 3 to 4 log increase in reporter expression was observed. We also infected a A549 cell over-expressing CD209 (DC-SIGN). DC-SIGN is a C-type lectin receptor that has been shown to enhance influenza A virus infection by binding to viral glycoproteins and acting as an endocytic entry receptor [16]. We observed enhanced infection of the A549(DC-SIGN) target cells by Ha-IAV and Ha-IBV pseudoviruses. However, such enhancement was not observed in the infection of Ha-IDV pseudovirus (**Figure 2C**). The mechanism for this discrepancy is unknown, but could be related to differences between IAV and IBV HA and IDV HEF.

Influenza A viruses contain the hemagglutinin (HA) surface glycoprotein, which can bind to sialic acid receptors on red blood cells (RBCs), causing hemagglutination. Influenza virus-mediated hemagglutination assay is a classic tool to quantify virus concentration and to measure specific antibody responses [17]. To confirm that the Ha-IAV particles assembled contain hemagglutinin and can functionally cause hemagglutination, we performed hemagglutination assays using turkey red blood cells resuspended in PBS and mixed with serially diluted Ha-IAV(A/Texas/37/2024-H5N1) particles. As shown in **Figure 3A**, we observed the hemagglutination of RBCs using undiluted viral particles and particles diluted to 1:2 and 1:4. We also observed partial hemagglutination from particles diluted to 1:8, but no hemagglutination at higher dilutions (1:16 to 1:512). These results demonstrated that Ha-IAV particles possess similar properties as wild-type influenza A virus in causing hemagglutination.

**Figure 3.**
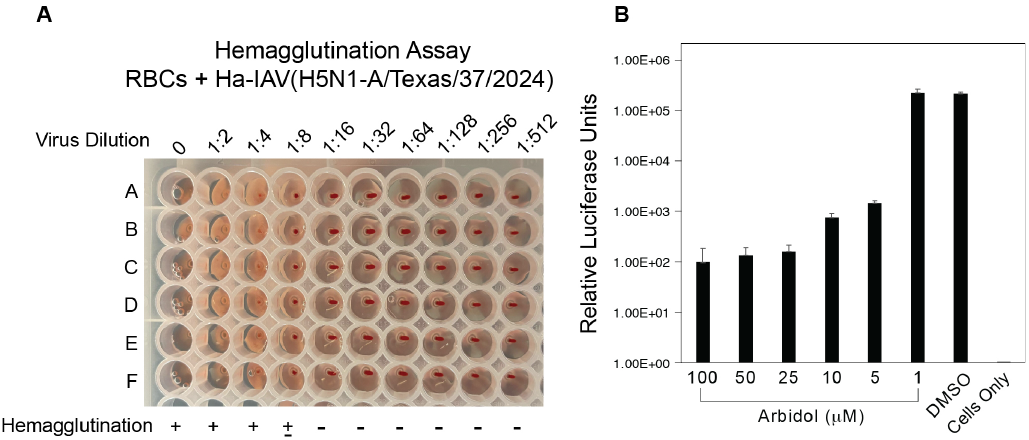
Validation of HA function in Ha-IAV. (**A**) Ha-IAV(A/Texas/37/2024-H5N1) pseudovirus-mediated hemagglutination reaction. Turkey red blood cells were mixed with serially diluted pseudoviral particles, incubated for 30-60 minutes, and observed for hemagglutination. (**B**) Arbidol inhibition of Ha-IAV(A/Texas/37/2024-H5N1)(Luc) infection. Cells were pre-treated with DMSO or with Arbidol at concentrations from 1 μM to 100 μM, and then infected with Ha-IAV(A/Texas/37/2024-H5N1)(Luc) for 18 hours. Luc expression was quantified. Luciferase assays were performed in technical triplicates, and the mean and standard deviation (SD) are shown.

HA-mediated influenza A entry is sensitive to entry inhibitors such as Arbidol, which is a small-molecule antiviral molecule blocking influenza A viral entry by binding and stabilizing the prefusion HA, inhibiting its conformational changes at low pH [18]. We tested the sensitivity of Ha-IAV(A/Texas/37/2024-H5N1)(Luc) to Arbidol inhibition, and observed a dosage-dependent inhibition of viral infection (**Figure 3B**).

### 3.3. Application of Ha-IAV(A/Texas/37/2024-H5N1) for quantifying neutralizing antibodies

To validate Ha-IAV(A/Texas/37/2024-H5N1) for rapid quantification of neutralizing antibodies, we tested a goat polyclonal H5N1 antiserum NR-10274, which was produced by immunization of a goat with a plasmid encoding the HA gene from N5N1 A/Vietnam/1203/2004 [19] followed by immunization with baculovirus-expressed HA proteins from both H5N1 A/Vietnam/1203/2004 and N5N1 A/Hong Kong/213/2003 [20]. The antiserum was serially diluted (1:5 dilution) and then incubated with Ha-IAV(A/Texas/37/2024-H5N1) (luciferase) pseudovirus. The antibody-virus complex or a control non-neutralized Ha-IAV(A/Texas/37/2024-H5N1) were used to infect cells for 18 hours for Luc expression. As shown in **Figure 4A**, we observed antibody concentration-dependent inhibition of Ha-IAV(A/Texas/37/2024-H5N1), and the IC50 (half maximal inhibitory concentration) was determined to be at 1:255.87 dilution (**Figure 4B**). These results demonstrate the application of the Ha-IVA(A/Texas/37/2024-H5N1) pseudovirus for rapid quantifying neutralizing antibodies.

**Figure 4.**
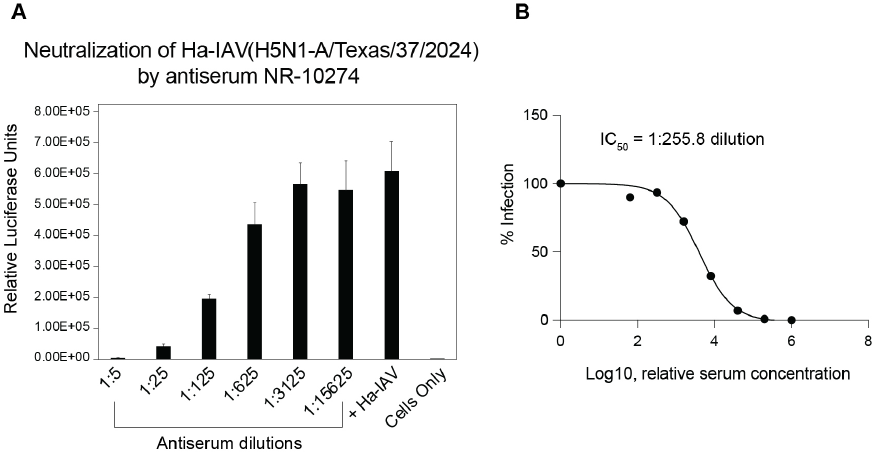
Rapid quantification of a neutralizing antibody using Ha-IAV(A/Texas/37/2024-H5N1). (**A**) Quantification of a standard neutralizing antibody with the Ha-IAV(A/Texas/37/2024-H5N1) particles. A H5 HA vaccinated goat polyclonal serum (NR-10274) was serially diluted and incubated with Ha-IVA for 10 minutes at 37°C, and then used to infect HEK293T cells. Neutralization activities were quantified by luciferase assay at 16 hours post infection. (**B**) The IC50 and neutralization curve were calculated using GraphPad PRISM software.

### 3.4. Validation of Ha-IVA(H5N1-A/Texas/37/2024) with infected cow antisera

Based on the results described above, we performed additional validation of Ha-IAV(A/Texas/37/2024-H5N1) for neutralizing assays using sera from influenza A-infected cows (two uninfected, one from 7 days post-infection, and one from 10 days post-infection). For comparison, we also performed an independent quantification of these sera using a hemagglutinin inhibition assay (HIA). As shown in **Figure 5A and 5B**, the Ha-IAV(A/Texas/37/2024-H5N1)-based neutralizing assay detected a strong neutralizing activity in the day 10 serum (IC_50_ 1:1102.5), while it detected minimal neutralization activity in the negative sera (IC_50_ dilutions 1:20.5 and 1:11.13) and low neutralization activity in the day 7 serum (IC_50_ 1:50.85). For comparison, the hemagglutinin inhibition assay also detected a strong inhibition in the day 10 serum (1:320), while it detected minimal antibody inhibition in the two uninfected controls and the day 7 post-infection serum (< 1:10). Assuming a protective threshold of 1:40 in the HIA assay, both assays identified the same group of positive and negative antisera. Nevertheless, the small size of the study only provides a proof of concept, but does not allow a direct correlation between the two assays.

**Figure 5.**
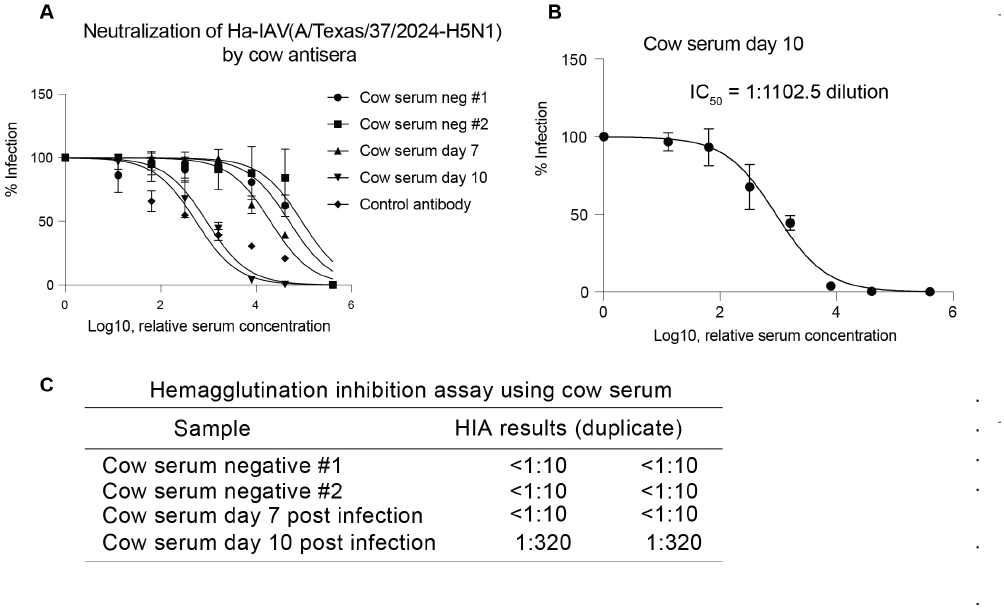
Proof-of-concept correlation study of Ha-IAV(A/Texas/37/2024-H5N1)(Luc) with hemagglutinin inhibition assay (HIA). (**A**) Results from Ha-IAV(A/Texas/37/2024-H5N1)(Luc)-based neutralization assay. (**B**) The IC_50_ and neutralization curve for day 10 cow serum were calculated using GraphPad PRISM software. (**C**) Results from hemagglutination inhibition assay.

## 4. Discussion

In this article, we describe the development and validation of a novel hybrid alphavirus-influenza pseudovirus that differs significantly from previous influenza pseudoviruses. Ha-IV offers a unique platform with several advantages for quantifying influenza virus types and subtypes. Structurally, Ha-IV is similar to a virus-like particle (VLP) and composed of major influenza virus structural proteins including HA, NA, M1, M2, and NP. Other pseudoviruses such as the VSV-or lenti-based influenza pseudoviruses only contain the influenza HA on the surface, and the other structural proteins are VSV or lentiviral structural proteins. The higher structural fidelity of Ha-IV is critical because it more accurately mimics the behavior of wild-type influenza viruses in drug screening, vaccine evaluation, and receptor binding studies; the structural protein on the surface of the pseudovirus must precisely match the conformational structure of the native pathogen [21]. Secondly, Ha-IV incorporates a reporter genome from the alphavirus replicon, which is known for its high efficiency and speed in gene expression. For example, in Sindbis virus alphaviral vector transduction, over 10^6^ CAT (chloramphenicol acetyltransferase) reporter molecules are produced within 7 hours [22], and 10^8^ CAT molecules per cell are accumulated within 16 - 20 hours post infection [23]. In addition, it has also been shown that transcription from the subgenomic RNA promoters occurs rapidly within hours, and the viral plus-RNA levels accumulate to 200,000 copies per cell [11, 12]. In our study, the Ha-IAV(H5N1-A/Texas/37/2024)-mediated Luc reporter expression can be detected as early as 4 hours post infection (**Figure 2**).

The use of rapid pseudoviruses can significantly enhance our ability to predict the impacts of HA/NA mutations and reassortments on the human-to-human transmission of influenza virus strains. Given that the highly pathogenic strains of influenza are BSL-3 level viruses, research on these strains is restricted to high-containment laboratories, limiting large-scale and timely evaluations of evolving viruses for their infectivity, host range, and responses to neutralizing antibodies. Our rapid Ha-IV pseudovirus platform allows virological and immunological assays to be performed in BSL-2 conditions within hours.

The classic method of measuring antibody neutralization is the hemagglutinin inhibition assay (HIA), which provides a quantification of antibody binding to HA. However, HIA has limitations in quantifying viral infectivity and neutralizing antibodies, especially given the variations of influenza subtypes in causing hemagglutination. For example, the H5 viruses inherently have a varied ability to cause agglutination compared to the H1 strains. The use of Ha-IV pseudovirus offers several advantages over classical HIA. First, Ha-IV pseudoviruses enable the direct measurement of viral entry mediated by influenza surface glycoprotein binding to receptors in a cell-based context, providing greater sensitivity and a broader dynamic range than HIA, which can be influenced by species-specific receptor differences. Second, unlike HIA, Ha-IV pseudoviruses are not limited to viruses that efficiently agglutinate erythrocytes and can be readily adapted to emerging strains or divergent influenza types. In addition, Ha-IV pseudoviruses are more amenable than HIA to high-throughput and quantitative analyses of neutralizing antibody responses.

In this study, we developed the first Ha-IDV pseudovirus. Influenza D virus (IDV) is a recently identified member of the family Orthomyxoviridae [6]. Since the initial discovery of the virus in cattle and swine, serological evidence of IDV infection has since been detected in a wide range of domestic and wild animals, which somehow resembles HPAI H5N1, highlighting its broad host range [7]. Despite this expanding evidence of cross-species transmission, IDV remains the least studied influenza virus, and its epidemiology and zoonotic potential are still poorly understood. Our Ha-IDV pseudovirus is expected to provide an important new tool in IDV research by providing a safe, flexible, and quantitative system to study viral entry and immune responses without handling replication-competent viruses. Because influenza D virus has limited reverse genetics tools and restricted availability of clinical isolates, Ha-IDV pseudovirus-based assays can fill a critical methodological gap and accelerate mechanistic, translational, and surveillance-oriented research on this emerging livestock-associated virus.

## 5. Patents

Patent applications related to the study have been filed.

## Author Contributions

Conceptualization, B.H., F.L. and Y.W.; methodology, B.H., F.L., D.W., K.K.H., and Y.W.; validation, B.H. and Y.W.; investigation, B.H., D.Y. and E.M., K.G., D.W.,; resources, B.H. and F.L.; data curation, B.H., K.K.H. and Y.W.; writing—original draft preparation, Y.W.; writing—review and editing, B.H., F.L., K.K.H., and Y.W.; visualization, B.H.; supervision, B.H. and Y.W.; project administration, B.H.. All authors have read and agreed to the published version of the manuscript.

## Data Availability Statement

results and data generated during this study are included in this article and are available upon request. All reagents generated in this study are available for research use with a Materials Transfer Agreement.

## Acknowledgments

We thank the Maxwell H. Gluck Equine Research Center, University of Kentucky for providing serum samples. The authors have reviewed and edited the output and take full responsibility for the content of this publication.

## Abbreviations

The following abbreviations are used in this manuscript:

Ha-IV: Hybrid alphavirus-influenza pseudovirus
HIA: Hemagglutinin inhibition assay
VSV: Vesicular stomatitis virus
CAT: Chloramphenicol acetyltransferase

